# Mismatch negativity develops in adolescence and independently of microglia

**DOI:** 10.1101/2025.10.14.682114

**Authors:** Anna M. Rader Groves, David A. Ricci, Joseph A. Wargo, Valentina J. Sutton, Hana S. Dalwai, Antanovia D. Ferrell, Bhoomi Desai, Jordan M. Ross, Connor G. Gallimore, Chloe L. West Jacobs, Georgia Bastos, Jessica L. Bolton, Fumiyasu Imai, Jordan P. Hamm

## Abstract

Higher brain functions and cognition undergo a critical period of development during adolescence, when psychiatric disorders such as schizophrenia typically onset. Understanding how developmental processes during adolescence interact with schizophrenia pathophysiology and risk remains a central goal in psychiatry. Here we show that a well-established biomarker of schizophrenia, mismatch negativity, matures during adolescence in mouse primary visual cortex, along with a strengthening of fronto-visual functional connectivity. Because microglia are implicated in schizophrenia risk and disease states, we further investigated what role microglia may play in the development of mismatch responses under physiological conditions. We found that microglial depletion with PLX5622 in adolescence arrests the development of resting oscillations in frontal areas, but does not affect the development of deviance detection, other signatures of visual context processing, or prefrontal–visual functional connectivity. Our findings suggest (a) a key component of mismatch negativity develops in adolescence, a period of vulnerability to schizophrenia, and (b) the development underlying this component does not require robust microglia activity, clarifying the developmental role of microglia in higher order visual processing.

## Introduction

Schizophrenia pathology follows a neurodevelopmental arc, and the symptoms typically onset in late adolescence and early adulthood^1^. An understanding of brain functions that both i) are altered in the disorder and ii) mature during adolescence could provide key pathophysiological insights. One key function altered in schizophrenia is sensory processing and integration^2,3^. The electrophysiological marker known as mismatch negativity (MMN) indexes these functions and is reliably deficient in schizophrenia in both visual and auditory domains^4–6^. MMN is most often measured during a sensory “oddball” sequence, wherein a series of repeated images is presented and occasionally interrupted by a different, “deviant” image; the difference in responses to these two stimuli constitutes the MMN^7–9^. Thus, MMN reflects how sensory cortex processes inputs in context and is thereby considered an index of integrative, sensory-cognitive function^10^. Consistent with this interpretation, MMN involves both sensory and frontal cortices, drivers of feedforward and feedback activity, respectively^11,12^. While MMN deficits are well-established in schizophrenia, and MMN function relies on regions that develop in adolescence (i.e., frontal cortex^13^), whether MMN develops during adolescence remains unclear^14–16^.

One factor contributing to this lack of clarity arises from the way in which MMN is commonly computed. Typically, researchers subtract EEG responses to deviant (“oddball”) from responses to redundant (“standard”) stimuli, which standardizes the measure but conflates two processes: *stimulus-specific adaptation* (reduced responses to repetition) and genuine *deviance detection* (enhanced responses to unexpected events)^17^. Stimulus-specific adaptation is present even in precortical processing stages, whereas deviance detection arises in cortex and requires both feedback from higher areas and cortical interneuron populations^7,18–21^. Importantly, when human studies have included the probability-matched control sequences to separate these components, it has been clear that deviance detection—not adaptation—is the MMN component reduced in schizophrenia^22,23^.

Given the role of fronto-sensory feedback in deviance detection^11,21,24–26^ and the protracted development of regions like prefrontal cortex over adolescence^13^, we hypothesize that visual cortical deviance detection matures in this time period, despite the common precept that V1 finishes developing prior to adolescence^27,28^. If confirmed, deviance detection could represent a precise measure of the developmental pathophysiology of this disorder and pave the way for the development of precision interventions in the prodromal phase.

Microglia – the brain’s immune cells – have come into focus in schizophrenia, as imaging^30^, postmortem^31^, and genetic studies^32^ point to aberrant activation and microglia-neuron interactions, but the field nevertheless lacks consensus as to the consistency of these observations in patient samples and, ultimately, the extent to which microglia contribute to schizophrenia pathology^33^. One hypothesis is that aberrant microglial activity during development contributes to schizophrenia neuropathology by overpruning synapses^34^, yet some counter examples challenge this model, suggesting a more general contribution to inflammation, rather than synaptic modifications^35^.

Even in the absence of pathology, the role of microglia in developmental synaptic pruning and circuit development in the cortex remains unresolved^36^, which ultimately limits advancements in neuropsychiatry. Microglia have been positioned as architects of developing neural circuits^37^. However, recent studies demonstrating limited impacts of removing microglia during development, specifically in cortex, have challenged the blanket application of this role across regions, ages, and functions^36,38^. These discrepant results call for a more careful approach to assessing how microglia impact the development of specific circuit elements. In particular, whether microglia are necessary for the maturation of higher order sensory processing and translational biomarkers such as MMN remains an important question.

Here we tested i) whether distinct components of the schizophrenia biomarker MMN develop in adolescence and ii) clarify the extent to which microglial cells contribute. We focused on the visual domain because visual perception and physiology are altered early in schizophrenia^2,3,39^ and visual mismatch responses are reduced in schizophrenia^6,40^. Moreover, recent work characterizing the role of microglia in visual development presents a gap in understanding the role of microglia later in visual development^38^. We show that visual deviance detection develops in adolescence in mice, along with functional connectivity between visual and prefrontal cortical areas. Stimulus specific adaptation, on the other hand, was present prior to adolescence. We further showed that depleting microglia (via dietary supplementation of PLX5622) in mid-adolescence (postnatal day (p)42-p50), but not peri-adolescence (p28-p36) or adulthood (p84-p92), augmented delta power and decreased beta power in prefrontal region ACa as measured in adulthood, aligning with past work and showing that microglia are needed for certain aspects of prefrontal circuit development. However, microglia depletion did not impact fronto-visual connectivity or, importantly, deviance detection. Together, this suggests that microglia may play a limited role in governing cortical development in adolescence – particularly with regard to the schizophrenia electrophysiological biomarker MMN.

## Results

### Deviance detection and fronto-visual functional connectivity develop over adolescence

To study how fronto-visual circuits and components of visual MMN develop over adolescence, we carried out *in vivo* electrophysiology in awake mice via chronic bipolar electrodes implanted in stereotaxically defined primary visual cortex (V1) and anterior cingulate area (ACa; **Fig. 1A**)--a mouse medial prefrontal region in the mouse that sends dense feedback connections to V1 that are necessary for visual MMN^20,26,41^. Mice were headfixed on a treadmill and presented with full-field oriented grating stimuli (100% contrast, drifting at 2 cycles per second for 500ms each, interleaved with a gray background inter-stimulus interval for 500-550ms; **Fig. 1A,B**) appearing in three separate, ≈5 minute sequences: a) an oddball run, wherein one orientation (e.g., 45 or 135 degrees) was presented repeatedly at 0.875 probability (the “redundant”), with rare orthogonally oriented “deviant” stimuli (e.g., 135 or 45 degrees, respectively) presented at 0.125 probability, b) an oddball flip run, where the prior redundant became the deviant, and vice versa, and c) a control run, where stimuli of eight orientations (0, 22.5, 45, 67.5, 90, 112.5, 135, or 155.5 degrees) were presented in a random order, such that all orientations appeared at .125 probability. Thus, across 3 runs, each orientation was presented in a context where it was a) redundant, b) deviant, or c) neither redundant nor contextually deviant (**Fig. 1B**). Our main analyses focused on deviance detection, which is captured by comparing the brain responses to context (b) versus (c).

**Figure 1:**
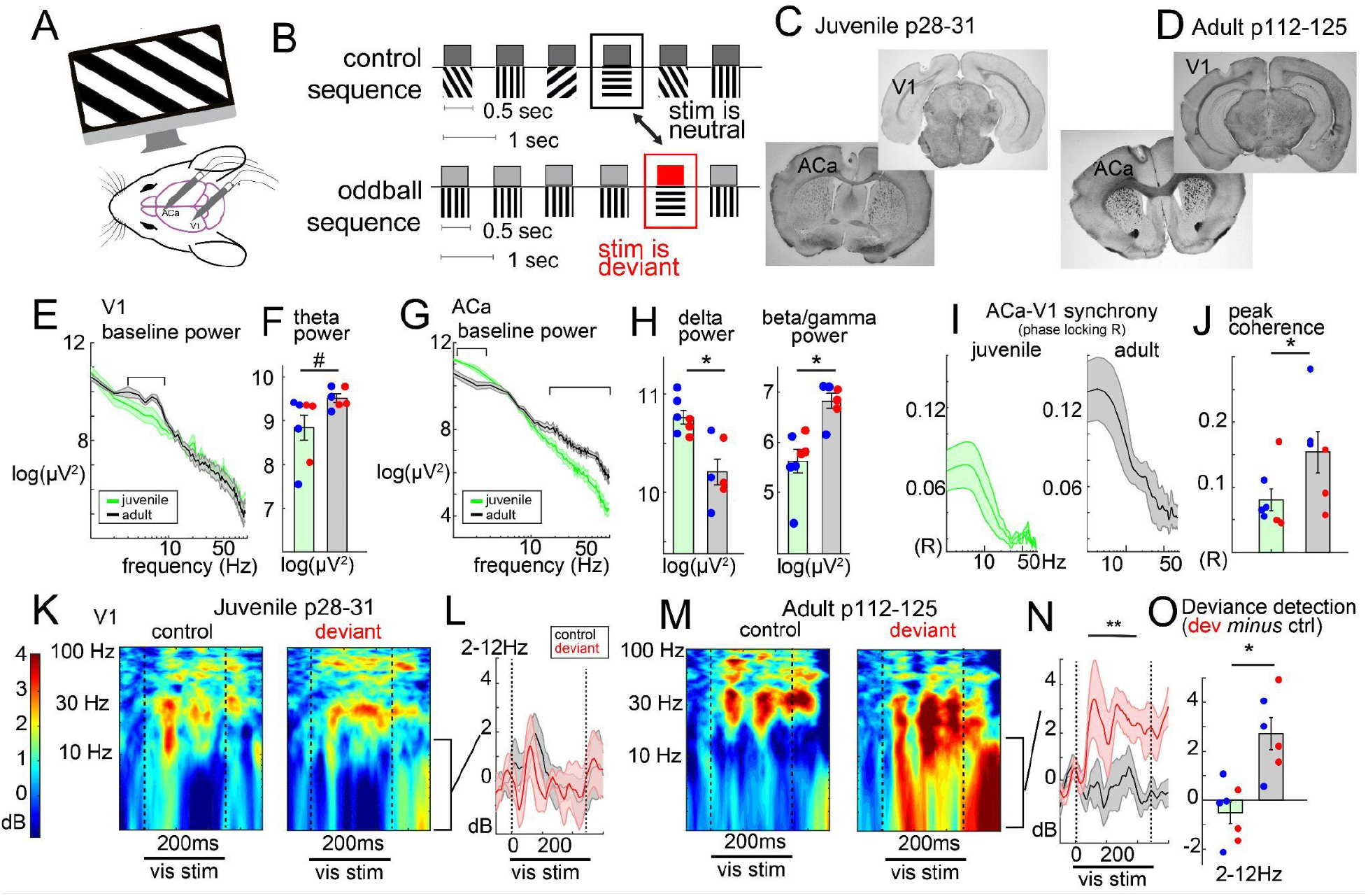
Frontal cortical circuitry and visual context processing develop in adolescence. A) Mice viewed full-field visual grating stimuli while local field potentials were recorded in ACa and V1. B) Visual stimuli were presented in many-standards control (top) and oddball (bottom) sequences to isolate contextual influences on visual processing, including deviance detection. C-D) Example histological verification of V1 and ACa electrode placement in (C) peri-adolescent and (D) adult mice. E) Resting power spectra and F) bar plots reflecting theta-peak power (6-8Hz) across male (blue) and female (red) mice. G-H) Same as E-F for ACa delta and beta/gamma power (averaged from 25-100Hz). I) ACa-V1 phase-synchrony spectra (phase lag coherence, *R*) during visual stimulation, averaged across control and oddball paradigms. J) Peak coherence values for each mouse (blue and red indicate male and female mice, respectively). K) Peri-adolescent stimulus-induced power (baseline adjusted as decibels) from V1 to the same stimulus orientations (45 or 135 degree) averaged across trials when it was contextually neutral control (left) versus contextually deviant. L) Averaged power from (K) for the delta/theta range (2-12Hz). M-N) Same as K-L in adult mice. O) Deviant minus control induced power from L, N, averaged from 100-400ms post stimulus for each mouse. *p<.05, **p<.01, two-tailed t-test

We studied juvenile (or pre-adolescent, p28-p31; n=7, 4 male) and adult mice (p112-p125; n=6, 3 male). At ≈p28, mouse V1 exhibits adult levels of basic feature selectivity (e.g. orientation, spatial frequency^42^), yet mice are just beginning -or have not begun-puberty^43^. Electrode locations were confirmed with post-hoc histology (**Fig. 1C,D)**. First, we analysed resting oscillatory power in V1 and ACa, i.e., in the ≈20 seconds prior to the start of visual stimulus presentation, during periods without locomotion. Across adolescence, resting power in ACa decreased in the delta range (**Fig. 1G,H**; 1-2Hz; F(1,9)=17.31, p=.002) and increased in the beta (13-25Hz; F(1,9)=9.13, p=.014), low gamma (25-55Hz; F(1,9)=12.95, p=.006), and high gamma range (65-100 Hz; F(1,9)=19.19, p=.002), consistent with past work^44^. Interestingly, peak theta power (6-8Hz) in V1 nominally appeared to increase over adolescence (**Fig. 1E**), but this only reached trend-level statistical significance and was not robust over mice (**Fig. 1F**, F(1,9)=4.07, p=.07). Juvenile and adult mice did not differ in the amount of time spent locomoting (**Fig. S1**, F(1,9)=1.98, p=.193).

During visual stimulation in adults, ACa and V1 exhibit phase coherence largely in the 3-12 Hz range that reflects bidirectional synchrony between these regions and, importantly, ACa’s role in supporting context processing and deviance detection in V1^26^. Here we find that this phase synchrony between ACa and V1 increased over adolescence (**Fig. 1G**, F(1,9)=5.90, p=.038). We also found that V1 deviance detection developed over this period (**Fig. 1K-O**). In adults, deviance detection was most prominent in the delta-theta band (2-12Hz; 100-400ms; t(6)=4.12, p=.009); **Fig. 1M-O**), as previously reported across species^8,19,26,45^, and this effect was absent in adolescent mice (t(7)=-1.18, p=.28; group by context interaction; F(1,9)=15.05, p=.004). These effects did not significantly interact with the sex of the mice, showing similar patterns in males and females. Interestingly, stimulus-specific adaptation (reduced responses to a stimulus in a redundant context relative to the same stimulus in a control context) was present prior to adolescence (**Fig. S2A**,**B**; 2-65Hz power, 100-400ms; t(6)=-2.99, p=.024) and did not differ between age groups (group by context interaction, F(1,9)=0.11, p=.739), demonstrating that it is not only mechanistically distinct from deviance detection, as previously shown^46^, but that it follows a separate developmental arc independent of late-developing cortical circuitry.

### Microglial depletion during adolescence alters prefrontal network oscillations in adulthood

Next we tested whether microglia were necessary for the adolescent development of frontal oscillations, fronto-visual connectivity, and deviance detection. We used a CSF-1 receptor inhibitor, PLX5622, mixed in Research Diets chow (1200ppm) to reversibly deplete microglia brain-wide during discrete periods in peri-adolescence (p28 to p36), mid-adolescence (p42-p50), adulthood (p84-p92), or not at all (control group; **Fig. 2A**). To control for the impacts of dietary changes, we switched food for all groups in each of the periods from their standard chow to the Research Diets chow, absent or present PLX5622. As expected, PLX5622 reduced microglial density in ACa and V1 by ≈90% after seven days of access to the PLX5622 chow, and then completely repopulated the cortex by one week after cessation of the drug (**Fig. 2B-D**).

**Figure 2:**
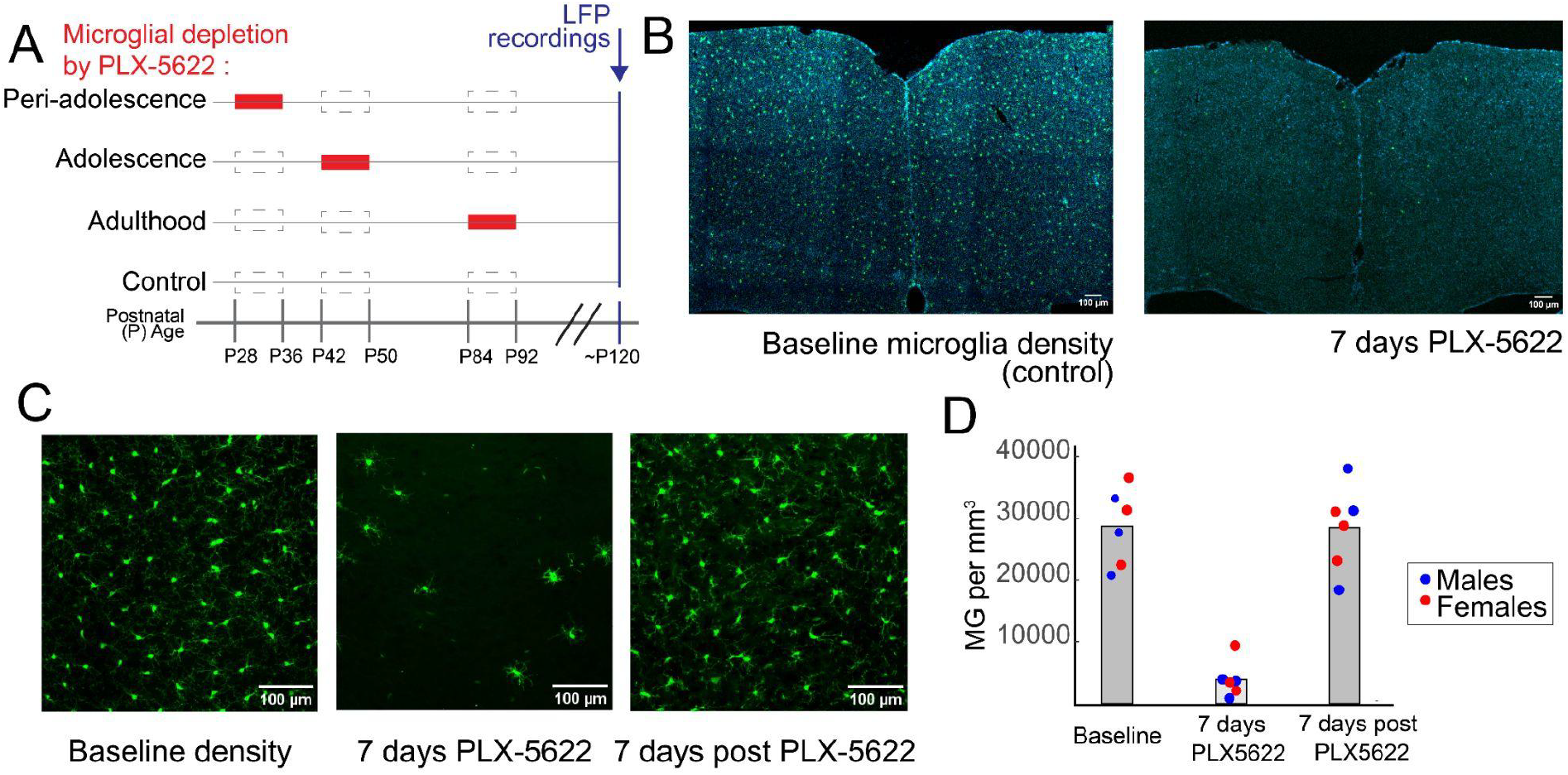
Microglia depletion via PLX5622 reversibly ablates microglia during discrete developmental periods. A) Schematic of the PLX5622 chow schedule across four groups: peri-adolescence (p28-26), adolescence (p42-50), adulthood (p84-92), and control (no PLX5622). Red boxes indicate periods of PLX5622 chow (i.e., experimental diet) and dashed borders indicate “pseudo-standard” chow (i.e., control diet). B) Microglia (labeled in CX3CR1-GFP mice, green) in coronal ACa (DAPI, blue) at baseline (left) and after 7 days of microglia depletion via PLX5622 (right). C) Microglia in ACa at higher magnification at baseline (left), at 7 days of microglia depletion via PLX5622 (middle), and at 7 days post PLX5622 cessation (right). D) Quantification of microglia depletion under conditions represented in (C).

First, in adult mice (p112-p125), we examined the impacts of prior microglia depletion on resting power, using the same methodology as above. We found that microglial depletion did not impact V1 resting power in any depletion group, for any frequency range (**Fig. 3A**).

**Figure 3:**
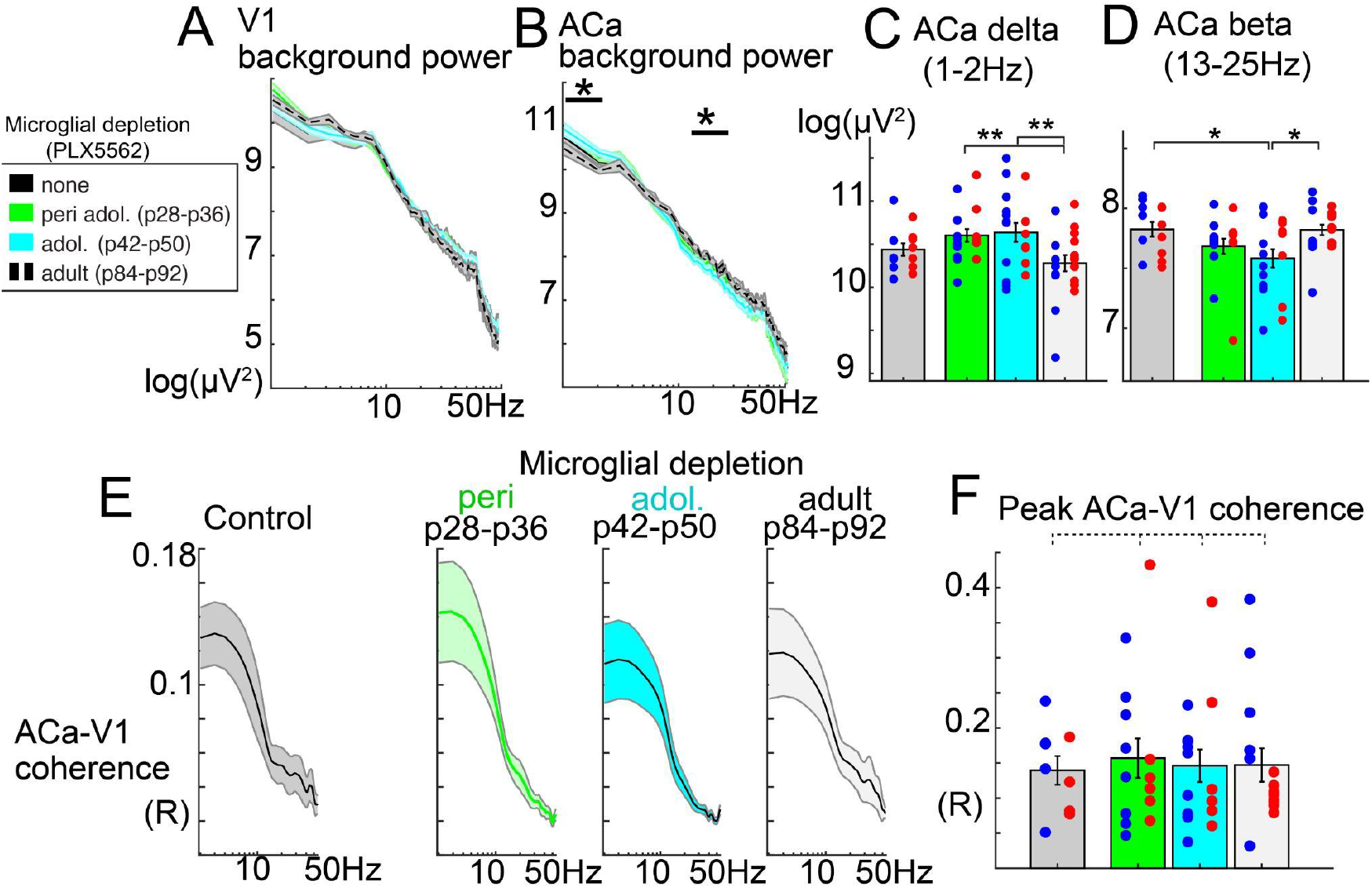
Developmental microglia depletion impacts local ACa oscillatory activity but not V1 or long-range ACa-V1 functional connectivity. All data reflect recordings from adult mice, p112-p125. A) Log-power spectra from V1 local field potential recordings during awake resting periods. B) Same as A in ACa. C) Group-wise barplots with overlaying scatters (each dot is one mouse; blue are males) for delta power and D) beta power from ACa. E) Phase coherence (inverse circular variance of phase lags) between ACa and V1, plotted across frequencies for each depletion group, and with F) peak-coherence values plotted across mice. Significance of group main effects (two-way ANOVAs between groups and sexes); *p<.05, **p<.01. Dotted lines indicate no significant group differences.

**Figure 4:**
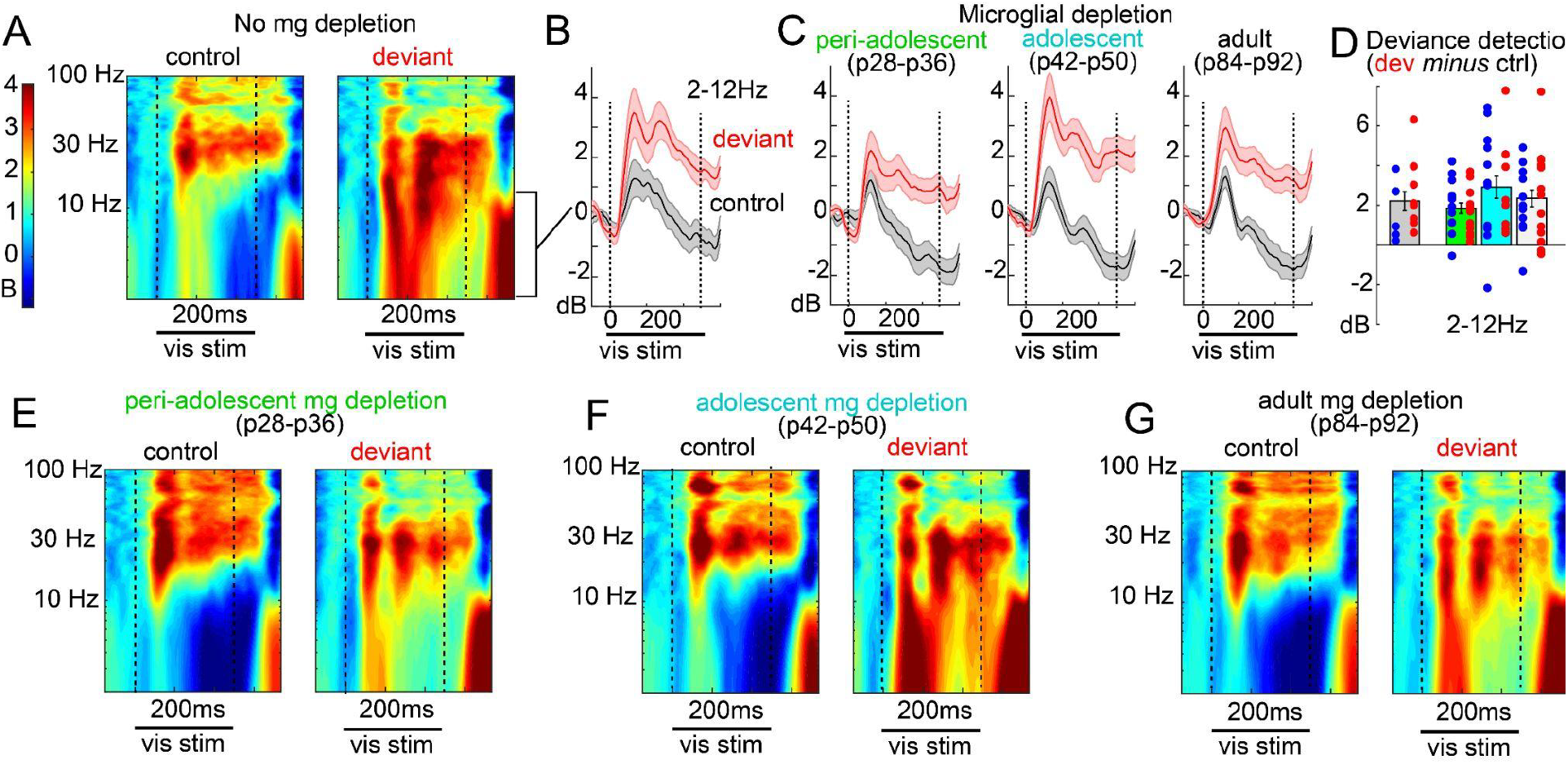
Developmental or adult-period microglial depletion does not impact deviance detection. A) Time-frequency spectra of local field potential recordings, averaged over mice, and averaged over trials when a given orientation (0, 45, 135, or 90 deg) was contextually neutral (control) or deviant evince B) clear deviance detection in the delta-theta range (line plots, averaged over mice) for the control group and C) all microglial depletion groups. D) Bar plot matching C, showing average delta-theta power from 100-400ms post stimulus onset. Dots are individual mice, and blue indicates male. E-G) Same as A for each depletion group.

However, in ACa, depleting during adolescence (p42-p50), and to a lesser extent, peri-adolescence (p28-p36), caused an increase in delta power (1-2Hz; group main effect; F(3,66)=3.65, p=.018) and a decrease in beta power (13-25Hz; F(3,66)=3.23; p=.029; **Fig. 3B-D**). Gamma-band power (26-100Hz) showed a similar trend as beta power in ACa, but did not reach statistical significance. These effects did not significantly interact with sex, and groups did not differ in the amount of time spent locomoting (**Fig. S1**, F(3,99)=0.974, p=.408).

Depletion of microglia in adulthood, after adolescence, did not impact V1 or ACa resting oscillations. Overall, this is consistent with the finding that frontal cortical (but not visual cortical) network-level oscillatory activity develops in adolescence (above) and that microglia play some role in this development^44^.

### Microglia are dispensable for the development of fronto-visual functional connectivity and visual deviance detection

While fronto-visual phase synchrony developed over adolescence (**Fig. 1**), microglial depletion did not impact this measure of long-range functional connectivity (**Fig. 3E,F**; peak-coherence value across spectrum, group main effect; F(3,53)=0.127; p=.944). Likewise, V1 deviance detection (which relies on feedback from ACa) also did not differ between depletion groups when measured in adulthood (2-12 Hz power, context by group interaction: F(3,73)=1.038, p=.381; **Fig 2A-G**), nor did stimulus-specific adaptation (**Fig. S2C**,**D**; context by group interaction: F(3,73)=1.262, p=.294). Given adult recordings were after p112 in all groups, while our PLX5662 treatment in the adolescent group stopped at p50 (and depletion was likely reversed within 3-7 days after that), the possibility that deviance detection develops in a period after ≈p57 (and nevertheless *is* dependent on microglia) could account for our null effects. We further examined deviance detection in a subset of p56 mice (n=7, 4 male), and found that, by this age, deviance detection is robust (**Fig. S3A;** t(6)=3.52,p=.013), and does not statistically differ from adults (context by age effect; F(1,9)=1.329, p=.279). Altogether, these results demonstrate that while some aspects of adolescent-developing ACa circuit activity, such as increased beta-band oscillations and decreasing delta oscillations, are arrested by microglial depletion, certain higher-order visual context processing functions such as deviance detection and long-range functional connectivity *also develop* throughout adolescence but do so even when the cortex is depleted of microglia by ≈90%.

## Discussion

Our results demonstrate that visual deviance detection, a key component of the mismatch negativity (MMN) biomarker, emerges in V1 over adolescence and is accompanied by increased functional connectivity between prefrontal and visual cortices. As visual MMN is reduced in schizophrenia^6^, the symptoms of which also emerge in late adolescence^1^, our work highlights the utility of this visual cortical biomarker in understanding core developmental disease processes. Moreover, these data add to a small but growing literature on the later development of higher order sensory functions, such as natural scene discrimination^29^, and invite a closer look at the maturation of primary sensory cortex beyond the critical window.

Past work in humans demonstrates that some form of MMN may be present across many stages of development including juveniles^14^. Here we clarify that deviance detection and stimulus-specific adaptation develop on distinct timescales, and emphasize the importance of experimental design that separates these components – especially in animal models. It warrants mention that the basic ability to detect and process visual novelty, at a *behavioral* level, is clearly present in young children (e.g., playing “peekaboo”), and past work has shown that later latency electrophysiological indices of deviance detection (e.g., the P300, occurring ≈100-150ms later than MMN) are present even in infants^47^. We show that deviance detection *in early visual cortex* (V1) emerges later, during adolescence, evident as early as 100 ms post-stimulus. This suggests that the ability to process context and prioritize novelty in early versus higher sensory regions continues to refine during adolescence, potentially reflecting the maturation of sensory filtering mechanisms that support attention and efficient perceptual processing^48^. Consistent with this notion, even the P300 shows a shortening of latency from childhood to adulthood^47^, suggesting a reverse propagation of predictive coding from high to low cortex across development. Our past work has shown that, in adult mice, deviance detection is present at multiple stages in the cortical hierarchy^18,49,50^. Future work should test this notion of reverse propagation in the visual system across development, with the prediction that deviance detection will be present in higher cortical areas earlier in development, along with behavioral indices of novelty detection.

Adolescence represents a critical window for the refinement of long-range cortical networks, as evidenced by increases in white matter continuing well into early adulthood^13,51^. The increase in ACa–V1 phase synchrony observed here suggests that enhanced oscillatory coordination may be a signature of this development in mouse models. The emergence of deviance detection during this same period suggests a functional link between structural and network-level maturation. Past work in rodents^20,26,52^ has demonstrated that this ACa-to-V1 feedback projection supports visual context processing (including deviance detection), consistent with human lesion studies showing a critical role of prefrontal regions in sensory context processing^53^. Other functions that this network is involved in, such as visuospatial attention, have also been shown to develop in adolescence as well, but involve beta/gamma band coordination between ACa and V1^54,55^ – a frequency band distinct from our 3-12Hz range reported here (Fig. 1) and elsewhere^26^. Whether this long range synchronization in these two frequency bands – delta/theta and beta/gamma – reflect distinct cellular mechanisms is an important question for future work.

We further examined whether microglia, posited to sculpt cortical circuitry in development^37,56,57^, are necessary for the development of visual context-processing functions. Transient microglial depletion during distinct developmental windows revealed selective effects on local frontal network activity – consistent with an arrest of typical development – but no impact on long-range synchrony or deviance detection. Specifically, mid-adolescent depletion increased delta and reduced beta oscillations in ACa when measured in adulthood, which matched the developmental changes in these bands seen in our typically-developing model (Fig. 1). This is also consistent with prior work implicating microglia in establishing mature patterns of frontal network activity^44^. However, the absence of effects on ACa–V1 coherence or visual MMN argues that the developmental emergence of context processing and top-down feedback is largely microglia-independent.

Mixed evidence for the role of microglia in postnatal circuit development^38,57–61^ highlights the importance of precisely defining the subprocesses and circuit elements under study, e.g., sensory context-processing as opposed to generally “cognition”. However, heterogeneous approaches^36^ – including the method of microglia depletion (e.g, the more cell-specific PLX-5622 versus the more commonly used PLX-3397)^62^, route of administration (e.g., acute delivery via surgery versus non-invasive systemic delivery brain-wide)^63^, duration of depletion, and definition of adolescence – all complicate the extrapolation or comparison of findings and, ultimately, preclude a generalized definition of the role of microglia in supporting cortical development. Our data adds to two previous studies that addressed the necessity of microglia in adolescent prefrontal cortical development^44,64^. These studies, which focus on prefrontal neuronal morphology and prefrontal-dependent behaviors, find microglia necessary in adolescence for the development of these functions. Here, we report that, where comparable measures were taken – specifically prefrontal oscillations – microglia in adolescence played a similar role but are dispensable for the development of fronto-visual circuit measures. Notably, Popplau and colleagues depleted microglia during peri-adolescence via PLX-3397, a less selective CSF-1R inhibitor than PLX-5662^62^, which could account for the broader effects observed. A similar study on mostly *pre-adolescent* V1 development used PLX-5662 (as in our study) and found much more limited contributions from microglia^38^.

The experimental design used here allowed us to deplete microglia reversibly in discrete periods without perturbing or stressing the animals through surgery or extensive handling. However, the several advantages of PLX-5622 chow (i.e., non-invasive, greater selectivity, reversible) required a few trade-offs that should be recognized in interpreting the results. Firstly, we chose not to deplete microglia in a specific region of interest, which would require surgery and generate an immune response, likely interfering with microglial function. However, this precludes us from assessing the necessity of microglia in a given region. Additionally, in order to specifically target discrete periods in development, we were unable to ensure complete elimination of *all* microglia (i.e., Csf1r^ΔFIRE/ΔFIRE^ mice^58^). Remaining microglia or other potential compensatory responses (e.g., astrocyte activity) would not reflect *typical* development of these processes; nonetheless, we cannot rule out that that a very small portion (∼10%) of microglia may have contributed to MMN and fronto-visual development in adolescence. Finally, we chose three timepoints for manipulation: peri-adolescence and (mid-) adolescence to assess the specificity of these periods for any developmental processes, and adulthood to control for any changes generated by the loss of microglia independent of developmental stage. So while the data presented here are specific to the role of microglia *during adolescence*, it is possible that microglia play an organizational role early in circuit development upon which later development scaffolds to generate mature MMN and fronto-visual circuitry.

Our data specifically addressed how microglia shape the development of sensory cortical neurophysiology and MMN, which is more specific to schizophrenia than prefrontal neuropathology^3,65,66^, and demonstrate that, in contrast to other cortical measures reported reliant on microglia, adolescent development of fronto-visual connectivity and MMN can proceed independently. This pattern of findings fits with the emerging picture of microglia as developmental regulators of circumscribed function (e.g., shaping prefrontal circuit properties)^36^. Whether these restricted impacts are due to region-specific developmental trajectories, function-specific redundancies, or altered compensatory responses requires further investigation.

In summary, this study identifies adolescence as a key period for the maturation of deviance detection and fronto-sensory connectivity, providing a mechanistic bridge between neurodevelopmental timing and a well-validated schizophrenia biomarker. Moreover, it demonstrates that while microglia influence local prefrontal network maturation, they are dispensable for the development of higher-order visual context processing. These findings highlight MMN as a tractable index of cortical circuit maturation and offer a refined framework for understanding how distinct developmental processes—neuronal, glial, and network-level—converge to shape neuropsychiatric risk.

## Methods

### Animals

All experimental procedures were carried out under the guidance and supervision of a) the Nathan S. Kline Institute for Psychiatric Research (NKI) Division of Laboratory Animal Resources or b) Georgia State University (GSU) Division of Animal Resources and in accordance with the approved Institutional Animal Care and Use Committee (IACUC) protocols at GSU and NKI. Male and female mice of C57BL/6 origin were used for all experiments. All adult mice (Figs. 1-4) were aged p112-125 and were of the CX3CR1-GFP strain. The peri-adolescent mice were C57BL/6 wildtype mice, and the P56 mice were GCaMP6-reporter mice. Cages were kept in a 12-hour light/dark cycle with one to five littermates of the same sex and depletion group (as applicable).

### Microglia Depletion via PLX5622

PLX5622 (Cayman Chemical), a CSF-1R inhibitor known to reversibly deplete microglia brainwide, was administered via chow during discrete time periods. All adult animals received eight days of non-standard chow during three time periods in development: peri-adolescence (p28-36), adolescence (p42-50), and adulthood (p84-92). Cages were randomly assigned to a depletion group and animals received chow with PLX5622 (AIN-76A Rodent Diet, D19101002 with 1200 ppm PLX5622, Research Diets) during the corresponding epoch. During the other two epochs, animals received a pseudo-standard chow matching the composition to the PLX5622 chow but lacking PLX5622 itself (AIN-76A Rodent Diet, D19101002, Research Diets). For example, a cage of animals assigned to adolescence depletion received the pseudo-standard chow from p28-36, the PLX5622 chow from p42-50, and the pseudo-standard chow from p84-92, with standard chow (Laboratory Rodent Diet, 5001, LabDiet) available ad libitum at all other times. Control animals received the pseudo-standard chow across all three periods.

### Surgery and Training

Electrode implantation and headplate fixation were carried out as previously described^26^. In short, mice were anesthetized using isoflurane (∼0.8-3%) and provided local and systemic peri-operative pain management. A titanium headplate was secured to the skull, then sterile bipolar electrodes (twisted together in pairs; Plastics One, Roanoke, VA, USA) were dipped in DiI and implanted in stereotaxically defined V1 and ACa unilaterally (left hemisphere). The second set of twisted contacts were grounded to the contralateral skull. Coordinates used for adult and juvenile are as follows: adult V1, X = 2 mm, Y = −2.92 mm, Z = dura; adult ACa, X = −0.35 mm, Y = +1.98 mm, Z = 0.9mm; juvenile V1, X = 1.8mm, Y = −2.6mm, Z = dura, juvenile ACa, X = −0.3mm, Y=1.8, Z = 0.8mm.

After surgery, mice received three days of head fixation training on a manual treadmill to improve comfortability and reduce stress. Training progressively increased in duration up to 45-60 minutes. During training, mice were exposed to the many-standards control visual sequence described below. After recording, brains were extracted and post-fixed overnight in 4% para-formaldehyde for post hoc electrode placement confirmation using DiI. All electrode placements were confirmed by two independent scorers under widefield (1.25x) magnification, referencing the Allen Brain Atlas.

### Visual Stimuli

Visual stimuli used were previously described in depth in (^26^). In short, mice were head-fixed on a freely moving manual treadmill, and full-field square-wave gratings (MATLAB Psychophysics Toolbox^67^) were displayed on a LCD monitor positioned 45 degrees relative to the animal’s head, approximately 15 cm in front of the right eye. Gratings were 100% contrast, at approximately 0.08 cycles per degree, drifting at 2.0 cycles per second to the right and/or upward, for 500ms. The interstimulus interval was a 500-550ms long gray screen of 50% luminance. Two types of visual sequences were presented. In the many-standards control sequence, gratings were randomly presented at eight orientations (0, 22.5, 45, 67.5, 90, 112.5, 135, or 155.5 degrees), each with equal likelihood of presentation. The oddball sequence comprises two orientations separated by 90 degrees. In the first half of the sequence, orientation A is selected as redundant (87.5% probability) and orientation B is selected as deviant (12.5% probability). In the second half, the orientations are “flip-flopped” such that orientation A becomes deviant and orientation B becomes redundant. We presented all mice with 6 runs: 2 oddball, 2 oddball flip, and 2 controls. The A/B pairs could be 0/90, 30/120, or 45/135. Sequences include 250 − 350 trials lasting 1.025 seconds in duration (500ms of drifting gratings with a 500-550ms blank gray screen in the interstimulus interval), totalling ≈4.5 minutes per sequence.

### Electrophysiology

During the presentation of visual stimuli, local field potentials were recorded via bipolar electrodes implanted in V1 and ACa. The electrodes were connected via insulated cables to a differential amplifier (high-pass: 0 Hz, low-pass: 500 Hz, gain: 1K, Warner instruments, DP-304A, Holliston, MA, USA). Electrical signals were then passed through a Digitmer D400, a 60Hz noise cancellation machine (Mains Noise Eliminator, Letchworth Garden City, UK) in order to adaptively remove line noise at 60Hz, avoiding waveform distortions that result from bandpass filters. Juvenile animals were ages p28 to p31 and adult animals were p112 to p125 at the time of recording. Locomotion was recorded via a photodarlington sensor attached to the treadmill (1kHz sampling) or by video as previous described^19,26^ (60 frames per second, Logitech C920 HD Pro webcam mounted ≈10 cm away from the mouse’s face, resampled to a binary locomotion trace). Juvenile, adult, and all depletion groups did not differ in the amount of time spent locomoting (Fig. S1). As this was typically ≈1% to 15% of the recording time, we excluded these data from all analyses and focused on time periods/trials without locomotion.

### Analysis

LFP data were manually inspected, and trials or segments (for resting state) with excessive signal (>≈5 std devs) in either V1 and ACa were excluded (between 0 and 50 across runs). All analyses focused on trials and segments where the mouse was not locomoting in the present or previous trial segment or trial. All trial numbers were matched between the number of retained trials in the comparison condition (deviant or redundant) and control run. For the deviant conditions, this was an average of 15 trials (between 11 and 21 across mice) for each orientation. For redundants, this was an average of 44 (between 30 and 50). Thus, we used separate control averages for the main deviance detection analysis (figure 1K,M) and for stimulus specific adaptation analyses (figure S2). For assessing deviance detection or stimulus specific adaptation in the juvenile vs adult analyses, we focused only on the 45/135 degree oddball runs, as these were recorded in all mice. For all other analyses and groups, we included both runs (which could have been 0/90 or 30/120deg). For quantifying the response to the control context, we additionally excluded trials where the orientation on the previous trial (≈500ms prior) was the same orientation or within 22.5 degrees of the current trial. This was done in order to control for short term adaptation effects that would affect the response to the control and lead to overestimates of deviance detection.

Once trials were selected, ongoing data were converted to the time-frequency domain with a modified morelet wavelet approach with 100 evenly spaced wavelets from 2-to 100-Hz, linearly increasing in length from 0.5 to 20 cycles per wavelet, applied every 10ms from 300ms pre-to 700ms post stimulus onset (200ms post-stimulus offset) as previously described^26^. Stimulus-induced power spectra (1-100 Hz) were computed for all three conditions (control, redundant, deviant) for each mouse and baseline-corrected by converting to decibels relative to the 100ms prior to stimulus onset, using EEGLAB^68^.

Interregional phase synchrony was quantified by taking the phase difference for each frequency (2-100 Hz) between ACa and V1 for all measurements from 100 ms pre-to 100ms post-stimulus offset and calculating the 1-circ variance (R-statistic) of these lags. We averaged these values across all available trials and conditions (as oddball and control runs did not vary systematically). For comparisons, we took the maximum coherence value from 2 to 55 Hz for each mouse. For the microglia depletion studies, there were 6 mice (evenly spread over groups) whose coherence spectra exceeded 6 standard deviations from their group means, suggesting artifactual coupling. They were removed from analyses, although this did not impact statistical conclusions.

For analysing resting power, data recorded prior to visual stimulation for each run was segmented in 1000ms locomotion-free, artifact-free epochs. Twenty epochs was the minimum available segments across mice, so used this for all comparisons in order to standardize signal/noise ratio. Segments were median normalized, and then converted to power spectra by taking squared absolute value of the output of the fast fourier transform of each segment, converted to log scale, and averaged over trials to yield a single spectrum for each mouse and region.

### Statistics

Only recordings from mice with histology-confirmed V1 targeting were used for V1 resting power and deviance detection analyses. Only recordings from mice with histology-confirmed ACa targeting were used for ACa resting power analyses. Only recordings from mice with histology-confirmed V1 and ACa targeting were used for coherence analyses. For the juvenile vs adult comparisons (figure 1), these numbers were the same for all analyses: n=7 (4 male) for juvenile; n=6 (3 male) for adult. For the microglial depletion experiments, these numbers varied, but exceeded 8 per group in all instances.

For resting power analyses, we focused on traditional frequency bands: 1-2Hz for delta; 3-9Hz for theta; 9-12Hz for alpha; 13-25Hz for beta; 25-55Hz for low gamma; 65-100Hz for high gamma. For power and coherence comparisons, we carried out factorial ANOVAs for each frequency band, using SEX and GROUP as between subjects variables. For deviance detection and stimulus specific adaptation analyses, we carried out mixed ANOVAs with CONTEXT as within subjects variables (control vs deviant; control vs redundant) and SEX and GROUP as between subjects variables.

## Acknowledgments

This work was funded by F31EY036279 (A.M.R.G.), R01EY033950 and R01MH128176 (J.P.H). Confocal imaging was supported by S10OD032336. The authors gratefully acknowledge the GSU Imaging Core for their assistance. The content is solely the responsibility of the authors and does not necessarily represent the official views of the National Institutes of Health.

## Notes

### Competing Interest Statement

The authors have declared no competing interest.

