## Supplement for "Mismatch negativity develops in adolescence and independently of microglia"

Supplemental Information

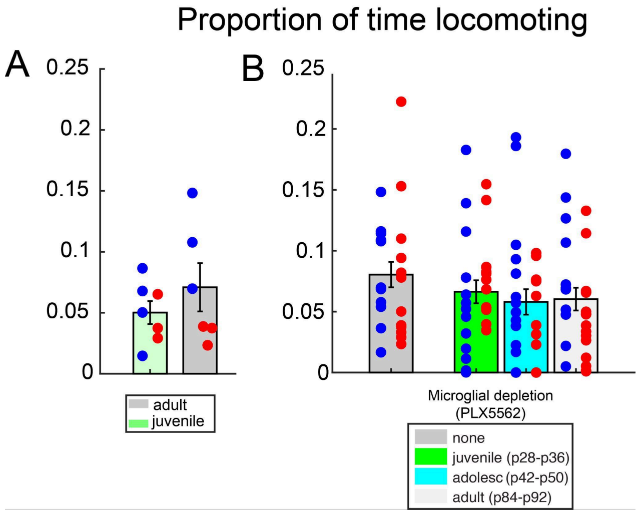


**Figure S1. Groups show similar locomotion during visual stimulation.** Locomotion was recorded via a rotary encoder or via video recording (60 frames per second). A) Average proportion of time that adult and juvenile mice exhibited locomotion during visual stimulation paradigms (control and oddball). Each mouse is a dot. Females are red and males are blue. B) Same as A but for microglial depletion groups. Total n for B reflects all mice recorded, which includes some mice that were excluded from main analyses based on histology (i.e. electrodes missed ACa or V1).

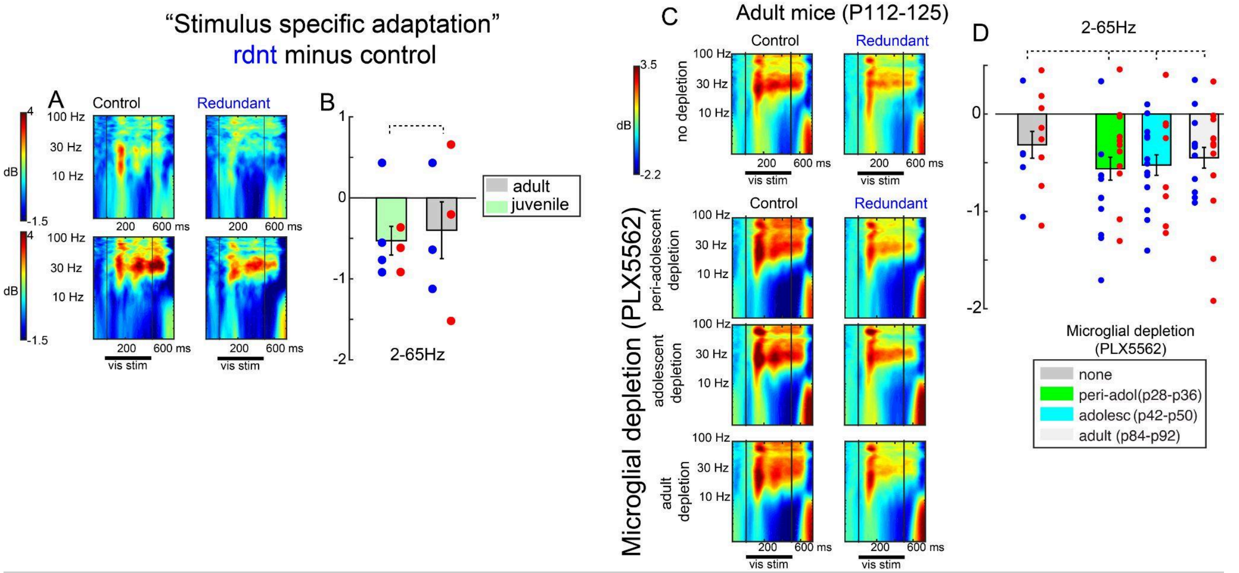


**Figure S2. Stimulus specific adaptation is present prior to adolescence and not impacted by microglia depletion**. A) Power spectra for juvenile (above) and adult mice to a stimulus when it was control vs redundant. Redundant average includes stimuli 3rd in the sequence or later, matched to the number of control trials (mean=44). B) bar plot, averaging power from 2 to 65Hz, from 100 to 400ms post stimulus onset. Each dot is one mouse, and males are blue. C,D) same as A/B, but for microglia depletion groups. Dotted lines indicate a lack of statistical differences among groups.

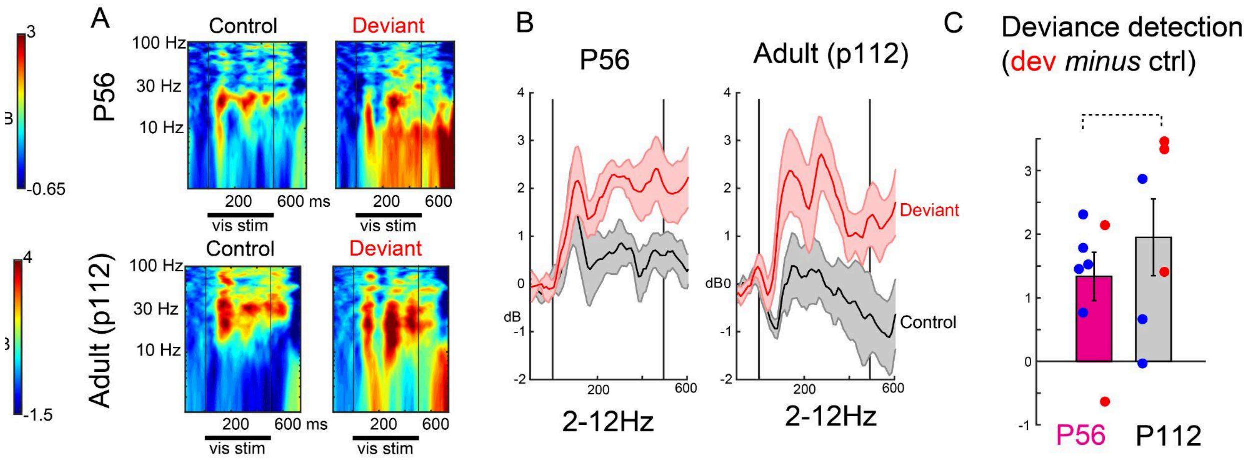


**Figure S3. Deviance detection is present at P56.** A) A) Power spectra for P56 (above) and adult mice to a stimulus when it was control vs redundant. Spectra averaged over 15 trials over 4 orientations (0, 45, 90, 135degrees). B) Trial averaged power from 2 to 12Hz. C) Trial-averaged 2-12Hz power averaged from 100 to 400ms post stimulus onset. Each mouse is one dot, males are blue. Dotted line indicates a lack of statistical differences between groups.
